# Targeting c-Myc with a novel Peptide Nuclear Delivery Device

**DOI:** 10.1101/2020.04.07.028977

**Authors:** Trinda Anne Ting, Alexandre Chaumet, Frederic Bard

**Affiliations:** Institute of Molecular and Cell Biology, Singapore 138673, Singapore; Department of Biochemistry, National University of Singapore, Singapore 119077, Singapore

**Keywords:** Toxin, delivery device, cargo, cancer cells, lymphoma, c-myc

## Abstract

Biologics such as peptides and antibodies are a well-established class of therapeutics. However, their intracellular delivery remains problematic. In particular, methods to efficiently inhibit intra-nuclear targets are lacking. We previously described that Pseudomonas Exotoxin A reaches the nucleoplasm via the endosomes-to-nucleus trafficking pathway. Here, we show that a non-toxic truncated form of PE can be coupled to peptides and efficiently reach the nucleoplasm. It can be used as a Peptide Nuclear Delivery Device (PNDD) to deliver polypeptidic cargos as large as Glutathione-S-transferase (GST) to the nucleus. PNDD1 is a fusion of PNDD to the c-myc inhibitor peptide H1. PNDD1 is able to inhibit c-Myc dependent transcription at nanomolar concentration. In contrast, H1 fused to various cell-penetrating peptides are active only in the micromolar range. PNDD1 attenuates cell proliferation and induces cell death in various tumor cell lines. In particular, several patient-derived Diffuse Large B-Cell Lymphomas cell lines die after exposure to PNDD1, while normal B-cells survive. Altogether, our data indicate that PNDD is a powerful tool to bring active cargo to the nucleus and PNDD1 could be the basis of a new therapy against lymphoma.

## Introduction

Biologics are a highly successful class of therapeutics that ranges from small peptides to large proteins such as antibodies. In contrast to small molecules, biologics can target proteins without hydrophobic pockets; they can disrupt complexes by interacting with specific domains. However, the inability of most biologics to cross the plasma membrane is a major limitation to develop therapeutics against intracellular targets. In particular, transcription factors, located in the nucleus, are largely inaccessible to most biologics.

Interestingly, cell surface receptors such as Epidermal Growth Factor Receptor (EGFR) and Low-density lipoprotein Receptor-related Protein 1 (LRP1, alias CD91), are able to translocate to the nucleus with their ligands ^1–3^. For instance, the bacterial toxin *Pseudomonas Exotoxin A* (PE) is a ligand for LRP1 and the related protein LRP1B^4^. PE is a 66 KDa protein comprising 3 domains: Domain I binds to the receptor LRP1, domain II has been described as a maturation domain and domain III contains an ADP-ribosylation domain that modifies the Elongation Factor 2 (EF-2) and inhibits host protein translation^5–6^. A 26-amino-acid peptide signal (PS) is also present at the N-terminus of the toxin and is cleaved before secretion in *Pseudomonas Aeruginosa*^5^. PE intoxication requires its transport via the retrograde pathway from endosomes, to the Golgi, then the ER before being translocated in the cytosol^7,8^. In addition, we recently showed that a fraction of PE is also delivered to the nucleus *via* endosomes that dock and fuse to the nuclear envelope shortly after internalisation. These recently-described endosomes are dubbed nuclear-associated endosomes (NAE)^2^. By removing the catalytic domain of PE, one can form a toxoid (non-toxic peptide) that could be targeted to the nucleus. Fusing this toxoid to other peptides could in theory allow their delivery to the nucleus. This approach could thus allow the specific targeting of nuclear transcription factors such as c-Myc.

C-myc is a proto-oncogenic transcription factor^9^. It is an elongated protein containing a basic helix-loop-helix (bHLH) and a leucine zipper (LZ)^10^. The LZ domains of both c-Myc and the transcription factor MAX interact to form a complex, afterwhich the combined bHLH domains binds to enhancer box sequences (E-BOX) of many pro-proliferation genes to drive their transcription^11,12^. Consistently, c-Myc is generally required for the growth of cancer cells^13^. Most genetic mutations of c-Myc are located in non-coding regulatory regions rather than in protein-coding regions, suggesting that dysregulation of c-Myc expression, rather than mutation, is the main driver of hyperproliferative cell growth^14^. Consistently, most cancer cells have elevated levels of myc and depend on this elevated expression^15,16^. For example, many B-cell lymphomas such as Burkitt’s Lymphoma and Diffuse Large B-Cell Lymphoma (DLBCL) depend on c-myc for pathogenesis^17,18^.

The loss of c-Myc can lead to tumor regression, proliferative arrest and/or apoptosis^19–20^. Although c-Myc is required in both normal and cancer cells to proliferate, cancer cells appear to be particularly sensitive to reduction of Myc levels, a phenomenon known as oncogene addiction^21^.

While c-Myc is an attractive target for cancer treatment, its nuclear localization and the lack of a hydrophobic pocket make it difficult to target by small molecules. c-Myc inhibitors have been developed: for example a small alpha-helix peptide able to inhibit the LZ interactions and called the H1 (S6A, F8A) peptide (hereafter called H1). H1 is a 14 amino acid (NELKRAFAALRDQI) competitive inhibitor derived from the interaction domain with MAX^22^. H1 reduces levels of c-Myc/MAX dimers and therefore inhibits transcription of downstream genes^22^. However, H1 is not membrane permeable, so cell-penetrating peptides (CPP) are employed for H1 intracellular delivery. However, the effective dosages of most CPP-H1 are in the micromolar range^23^, making it impractical for drug development. Another Myc inhibitor is Omomyc, a polypeptide that binds to the E-Box and competes with the c-Myc-Max dimer to inhibit transcription^24,25^.

In this paper, we describe the development of a truncated PE toxoid as a Peptide Nuclear Delivery Device (PNDD). We show that it is able to deliver various cargo to the nucleoplasm, including the H1 peptide. Using this approach, we observed an inhibition of c-Myc activity in various cancer cells, including lymphoma-derived cell lines. In many cell lines, the inhibitor induces a significant reduction in cancer cell proliferation and eventually inducing cell death.

## RESULTS

### PE domains Ia and II are sufficient for nuclear delivery

To determine the PE domains required to reach the nucleus, we generated four truncated toxoids devoid of the cytotoxic catalytic domain III. Here, PE amino acids are numbered from the first amino acid of the peptide signal. PE389 contains domains Ia and II, PE277 contains only domain Ia; PE212 and PE151 are truncated versions of domain Ia (Fig. 1A). We expressed these constructs in E.Coli under an Isopropyl *β* -D-1-thiogalactopyranoside (IPTG) inducible promoter. As most constructs were not readily soluble, we used urea extraction to obtain the recombinant proteins (Supplementary Fig. S1A-L). To examine nuclear trafficking of the constructs, we first compared their enrichment in NAE to that of wild-type (*wt*) PE. To visualise delivery to the NAE, we used MG63, human bone osteosarcoma cells expressing high levels of LRP1. The MG63 cells were incubated with the purified constructs labelled with HiLyte™ Fluor 488 dye (488 dye for short) together with *wt* PE labelled with HiLyte™ Fluor 594 dye (594 dye for short) for an hour. NAE were defined as PE-positive endosomes located above or below the nucleus, in close contact with the nuclear envelope as previously described^2^. We observed colocalization of PE389, PE277 and PE212 with *wt* PE (Fig. 1B). By contrast, PE151 signal was barely detectable.

**Figure 1.**
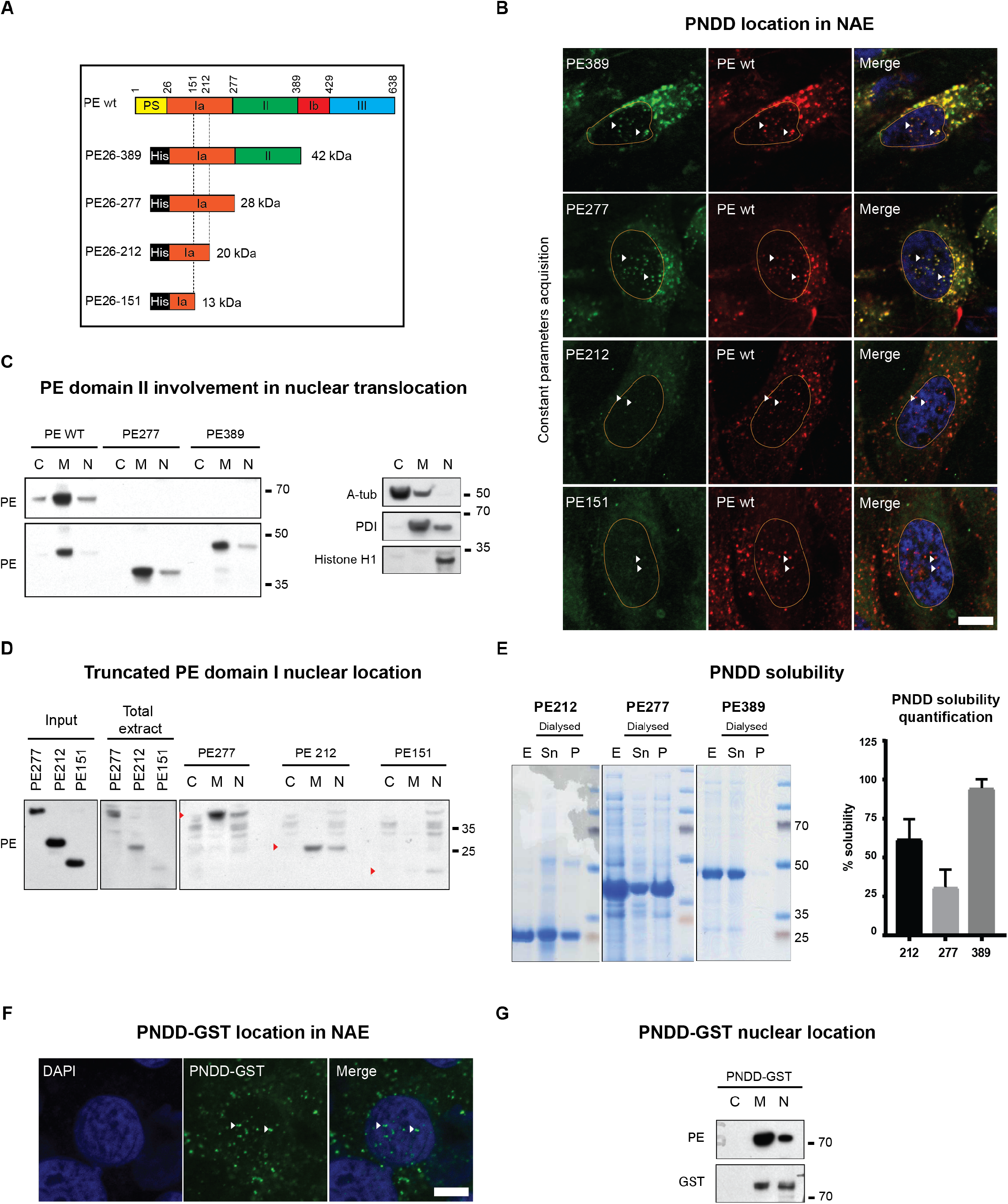
PE domains I and II are sufficient to carry cargo to the nucleus. (**A**) Primary structure of PE wt and different constructs tested for their ability to reach the nucleus. Name of the construct refers to amino acid numbers. PS: Peptide signal cleaved by *Pseudomonas Aeruginosa;* His: histidine tag for purification. Theoretical molecular weights are listed on the right. (**B**) Confocal Imaging of PE-HiLyte™ Fluor 594 dye and truncated PE-HiLyte™ Fluor 488 dye 1 hour after incubation on MG63 cells. White arrows show NAE structure containing both PE *wt* and truncated PE. Representative images of 3 independent experiments. Nucleus is delimited in orange. Images were taken at constant parameter acquisition settings (scale bar: 5 um). (**C**) Test of domain II involvement in truncated PE nuclear translocation. MG63 cell fractionation after 1 hour treatment with PE wt, PE389 or PE277 construct (left panel). Representative fractionation control: A-Tub, PDI and Histone H1 are used as fraction controls for respectively C: Cytosolic fraction; M: Membrane fraction; N: Nuclear fraction (right panel). Molecular weights are shown on the right. Representative images of 3 independent experiments. (**D**) Test of domain I sequence involved in truncated PE nuclear translocation. MG63 cell fractionation after 1 hour treatment with PE 277, PE212 or PE151 constructs. Total extract: cell lysate after 1 hour treatment; Input: purified protein; C: Cytosolic fraction; M: Membrane fraction; N: Nuclear fraction. Antibodies are labelled on the left of each blot. Molecular weights are shown on the right. Representative images of 3 independent experiments. (**E**) Coomassie staining showing PE389, PE277 and PE212 solubility after dialysis and centrifugation. E: Eluate before dialysis and centrifugation; Sn: supernatant containing soluble protein after centrifugation; P: Pellet containing insoluble protein after centrifugation. Molecular weights are shown on the right (left panel). Quantification of soluble protein proportion (right panel). Error bars at s.d. All purifications are shown in Supplementary Fig. S1. (**F**) Confocal Imaging of PNDD-GST-HiLyte™ Fluor 488 dye. Representative images of 3 independent experiments (scale: 5 um). (**G**) MG63 cell fractionation after 1 hour treatment with PNDD-GST. Molecular weights are shown on the right. Representative of 3 independent experiments.

We next checked the ability of the constructs to reach the nucleus by cell fractionation. We incubated MG63 cells with PE389 or PE277 for 1 hour before fractionation and found both constructs in the nuclear fraction, suggesting that domain II is not essential for nuclear translocation (Fig. 1C). PE212 appeared nearly as efficient as PE277 to be internalised and be enriched in the nuclear fraction (Fig. 1D). By contrast, PE151 was not detected in either total cell lysate nor nuclear fraction (Fig. 1D). In conclusion, PE389, PE277 and PE212 have comparable abilities to enter the nucleus, while PE151 appears to be poorly internalised. These results also suggest that domain II is not required for delivery in the nucleus.

To form an efficient delivery device, the toxoid selected must have good stability and solubility. Since the constructs are solubilised by urea for extraction, they need to be dialysed extensively before use, which can lead to aggregation and loss of material. We thus compared raw eluates, post-dialysis eluate supernatants and pellets by SDS-PAGE followed by Coomassie staining (Fig. 1E). The recovery of PE389 in the soluble dialysed fraction was 95%. By contrast, PE277 and PE212 only had 25% and 60% recovery respectively (Fig. 1E). Therefore, despite its larger molecular size, we selected PE389 to work with and hereafter dubbed it as Peptide Nuclear Delivery Device 0 (PNDD0).

To test the ability of PNDD to deliver protein cargo to the nucleus, we fused the 28 kDa Glutathione S-transferase (GST) in C-terminus of PNDD0. PNDD-GST was coupled with the 488 dye to study its subcellular distribution and found to localise in NAE after 1 hour of incubation with MG63 cells (Fig. 1F). PNDD-GST was also detected by both PE and GST antibodies in nuclear fraction after 1h incubation, suggesting that PNDD could deliver GST effectively to the nucleus (Fig. 1G).

### A chimeric cMyc inhibitor, PNDD1, is delivered to the nucleus

We next fused the c-Myc inhibitor H1 at the C-terminus of PNDD0, hereafter called PNDD1 (Fig. 2A). We found the solubility of PNDD1 to be similar to PNDD0, as seen from comparable levels of recovery after dialysis (Fig. 2B, S2A-B). We verified delivery to the nuclear fraction in MG63 cells after 1 hour exposure using cell fractionation (Supplementary Fig. S2C).

**Figure 2:**
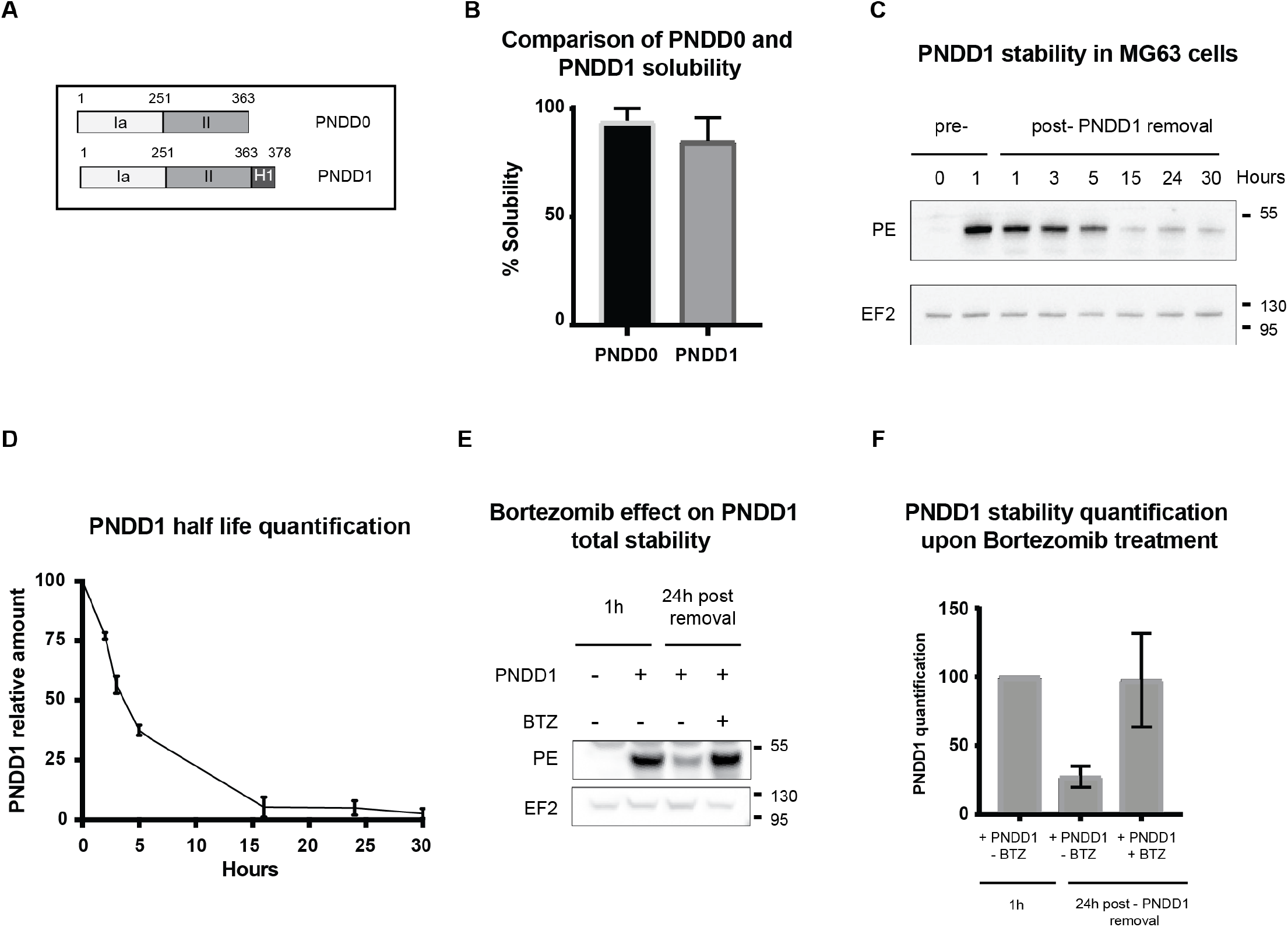
PNDD1 nuclear location and half life. (**A**) Schematic of PNDD0, PNDD1 (PNDD0 + H1 peptide). (**B**) Quantification of soluble PNDD1 proportion from Supplementary Fig. S2A-B and soluble PNDD0 proportions (Supplementary Fig. S1A-F). Error bars at s.d. (**C**) PNDD1 half-life: PNDD1 was incubated for 1 hour (pre-) on MG63 cells before being washed with PBS and replaced with culture media (post-). Cells lysates were obtained at different time points up to 30 hours and amount of intracellular PNDD1 was analysed by western blot to determine its half life in cells. EF2 was used as a loading control. Representative western blot of 3 independent experiments. Replicates are represented in Supplementary Fig. S2F. (**D**) Average quantification of PNDD1 bands in Fig. 2C and Supplementary Fig. S2F. Error bars at s.d. (**E**) Effect of proteasome inhibitor Bortezomib on PNDD1 stability. MG63 cells were incubated with PNDD1 during 1 hour with or without Bortezomib before wash, then harvested after 24h. MG63 cells were incubated for 1 hour with PNDD1 as a control for maximum uptake. EF2 was used as a loading control. Molecular weights are shown on the right. Triplicate experiments are represented in Supplementary Fig. S2G. (**F**) Quantification of PNDD1 in Fig 2E. Error bars at s.d.

To evaluate the fraction of internalised PNDD1 that reaches the nucleus, we quantified the ratio in the nucleus to total extract (Supplementary Fig. S2D) and using the nuclear located protein Max as a reference, we estimate that around 6% of internalised PNDD1 is found in the nuclear fraction after 1h of incubation. This seems consistent with the relatively small amount observed in NAE compared to the perinuclear pool of PE (Fig. 1B).

We next measured the kinetics of uptake and observed that cellular amounts of PNDD1 appear to saturate after one hour (Supplementary Fig. S2E). This result suggests that either the LRP1 receptor is saturated and/or degradative mechanisms start to balance out the uptake of additional molecules. To test for intracellular degradation, we pulsed MG63 cells with PNDD1 for 1 hour before incubation with normal media. Total cellular PNDD1 was quantified by western blot, leading to an estimated intracellular half-life of around 4 hours. An intracellular pool of PNDD1 can be detected as late as 30 hours post-wash (Fig. 2C-D, Supplementary Fig. S2F).

Next, we explored how PNDD1 is degraded. Since the toxoid is translocated in the nucleoplasm and the cytosol, its degradation is likely to be in part cytosolic. Treatment with the proteasome inhibitor Bortezomib largely increased PNDD1 stability (Fig. 2E-F, S2G). This suggests that proteasomal degradation is a major pathway and proteasome inhibition a potential way to increase the potency of PNDD1 in vivo.

### PNDD1 specifically inhibits c-myc transcriptional activity

To analyse whether PNDD1 is able to inhibit c-Myc, we established a Myc activity reporter cell line. We selected the epidermoid carcinoma cell line A431 because of their high c-Myc levels and ease of transfection. We introduced the firefly luciferase under the control of an E-BOX containing promoter. We also introduced a Cytomegalovirus (CMV) promoter-driven Renilla luciferase gene reporter as a control (A431/EBox-Luc/CMV-Ren cells, A431-mrep in short). In this system, inhibition of c-myc should lead to a specific reduction of the firefly luciferase signal (Fig. 3A).

**Figure 3:**
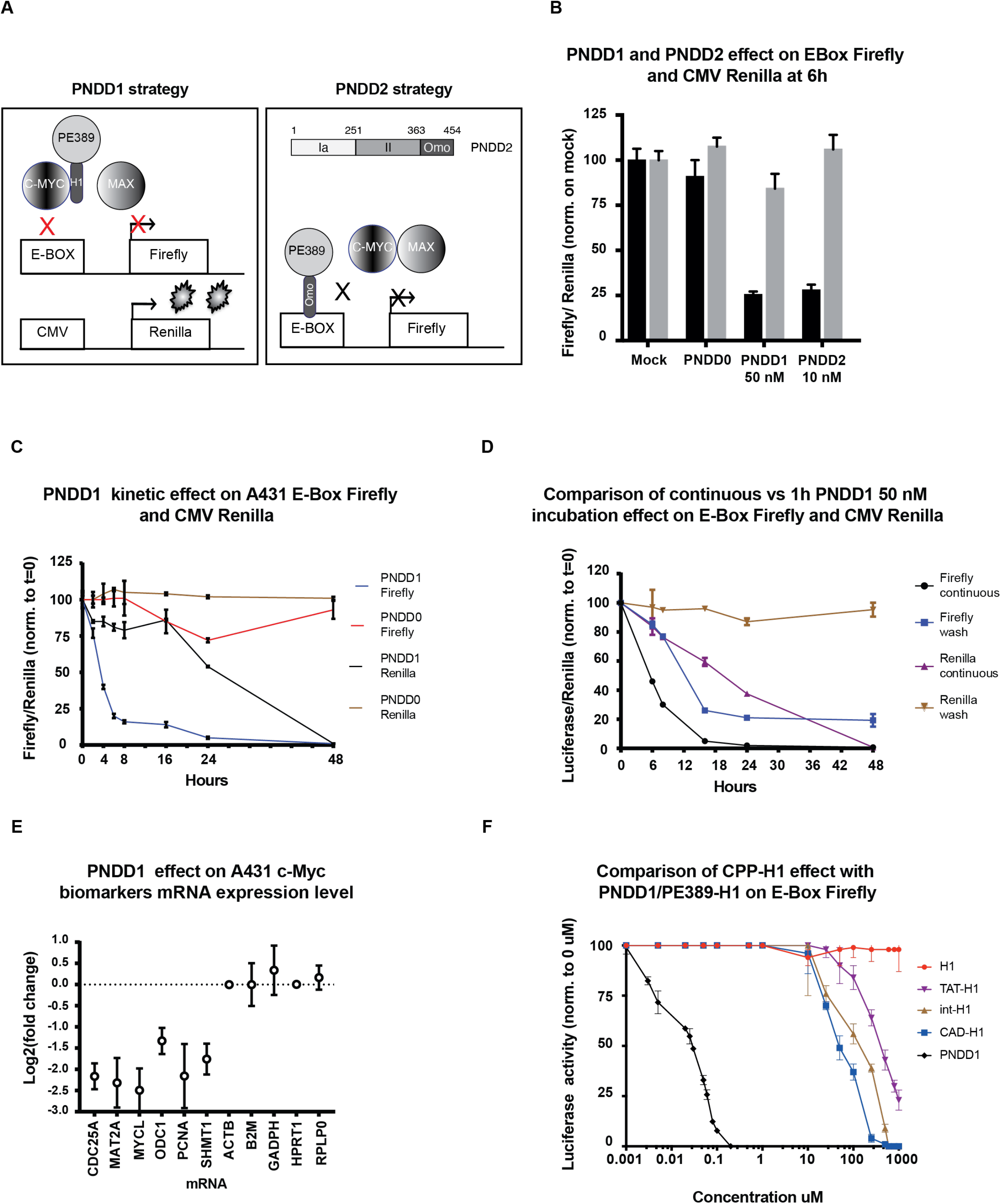
PNDD1 effect on E-Box luciferase. (**A**) Schematic of c-Myc reporter system in A431-mrep cells and mode of action of PNDD1/2. (**B**) PNDD1 and PNDD2 effect on c-myc reporter system. Firefly luciferase: black bars, Renilla luciferase: grey bars; A431-mrep cells were treated for 6 hours with 50 nM PNDD0, PNDD1 or 10 nM PNDD2. Error bars are s.d. (**C**) PNDD1 50 nM kinetics on c-Myc reporter system over 48 hours. (**D**) Comparison of continuous versus 1 hour PNDD1 50 nM incubation. (**E**) PNDD1 effect on c-Myc biomarkers mRNA expression level. A431 cells were treated with PNDD1 50nM overnight (16 hours) before RNA extraction. mRNA transcript levels of genes were quantified by RT-PCRQ and compared with or without PNDD1 treatment. Housekeeping mRNAs (HPRT1, GAPDH) whose expressions are not regulated by c-Myc are analysed in the same manner. Y axis shows the mRNA log2(fold change). Upregulated genes appear with negative log2(fold change) compared to housekeeping genes. Representative of 3 independent experiments. (**F**) Comparative dose response of cell targeting peptides (CPP) fused to H1 and PNDD0 (PE389) fused to H1 (PNDD1/PE389-H1) on A431-mrep cells after 6 hours treatment. X axis is shown in Log scale. Cadherin (CAD; LLIILLRRRIRKQAHAHSK;) EC_50_=75 mM, Antenapedia (Int; RQIKIWFQNRRMKWKK) EC_50_=200 mM and TAT (GRKKRRQRRRPPQ) EC_50_=500 mM. Error bars at s.d. Representative of 3 independent experiments.

We verified PNDD1 uptake and nuclear location in A431-mrep by cell fractionation after 1 hour treatment (Supplementary Fig. S3A). After a preliminary dose response evaluation, we determined that 25nM was an approximate EC50 for PNDD1 (Supplementary Fig. S3B). We next used 50nM to treat A431-mrep cells for 6 hours. In these conditions, Firefly luciferase activity was decreased by^~^75% while Renilla luciferase activity remained constant (Supplementary Fig. S3C). Treatment with 50nM PNDD0 did not have a significant effect on either reporters (Fig.3B and S3C).

Since our preliminary data with GST indicated that a larger peptide than H1 could be efficiently delivered by PNDD, we next tested coupling the 10KDa Omomyc to PNDD0 (hereafter referred to as PNDD2) (Fig. 3A). After verifying its cellular internalisation and nuclear enrichment (Supplementary Fig. S3D), PNDD2 was tested with the A431-mrep for 6 hours, leading to decreased Firefly luciferase activity in a dose dependent manner (Fig. 3B, S3E). Thus, the PNDD method of delivery is not limited to H1 but is adaptable to other Myc inhibitory cargos.

We next explored the kinetics of PNDD1 effects. A rapid 4-fold reduction of Firefly luciferase signal was observed within 6h hours, further decreasing to near undetectable activity by 24h (Fig. 3C). By contrast, Renilla luciferase activity was near normal up to 16h, after which it started to drop as well (Fig. 3C). We interpret this decrease as due to loss of cell fitness.

We next tested how a pulsatile exposure to PNDD1 would impact Myc activity. After 1h incubation with PNDD1, the supernatant was removed and cells washed. As expected Myc inhibition was less marked but appeared surprisingly lasting, between 24 and 48h (Fig. 3D, Supplementary Fig. S3F). Firefly luciferase activity rebound at longer time points, 72h and beyond (Supplementary Fig. S3F). This suggests that PNDD1’s effect on Myc activity is more prolonged than the half-life of PNDD1. By contrast with continuous exposure, Renilla activity was not significantly impacted by a 1h exposure (Fig. 3D).

Next, to validate the on-target activity of PNDD1, we measured the changes in mRNA expression of several c-Myc biomarkers by quantitative RT-PCR after exposure at 50 nM for 16 hours (Fig. 3E, S3G). Consistently, the expression of Myc targets CDC25A, MAT2A, MYCL, PCNA and SHMT1 were significantly down-regulated after PNDD1 treatment^26,27^. By contrast, mRNA levels of housekeeping genes are not affected (Fig. 3E, S3G).

Altogether, these results show that PNDD is able to deliver the H1 peptide and the Omomyc polypeptide to the nucleus, the fusion does not impair the inhibitory activity of the peptides and inhibition is rapid and lasting.

### PNDD1 is thousand times more potent than cell penetrating peptides

In order to compare the efficiency of PNDD with existing cell entry systems, we fused H1 with commonly used cell-penetrating peptides (CPP) such as HIV TAT (TAT), cadherin peptide (CAD) or antennapedia (Int)^28^. We performed dose response assays comparing these CPP-H1 and PNDD1 on A431-mrep after 6h incubation and at concentrations ranging from 1 nM to 1 uM (Fig. 3F, S3H). H1 peptide alone had no effect, suggesting an inability to penetrate the cell membrane. Among all the CPP-H1 fusion proteins, CAD-H1 was the most efficient with an EC50 of 75uM, followed by Int-H1 (EC50 = 200 uM) and TAT-H1 (EC50 = 500 uM) (Supplementary Fig. S3H). Some of these EC50 values are higher than previously reported^29^. This may be due to differences in peptide preparation and assay specificity. Nevertheless, in our hands, the EC50 of the most potent CPP fusion, CAD-H1 is^~^3000 times higher than PNDD1 (Fig. 3F, S3H, S3I).

### PNDD1 inhibits proliferation of different tumour cells

In order to test whether PNDD1 inhibition of c-Myc also affects cell proliferation, we tracked the growth of A431 cells for 8 days using time-lapse microscopy. A431 cell density was evaluated using phase contrast images of the culture and image analysis (Fig. 4A). After seeding at^~^10% confluency (5000 cells per well in a 96 well plate), the culture reached confluency by day 4 (Fig. 4A). When incubated with 50 nM of PNDD1, the rate of cell proliferation decreased, reaching only about 70% confluency at day 4. In addition, cell confluency started to decrease after 4 to 6 days, suggesting that cells started to die. The effect on cell proliferation was dose dependent, proliferation being slowed more with higher doses until completely blocked with 200nM PNDD1 (Fig. 4B).

**Figure 4:**
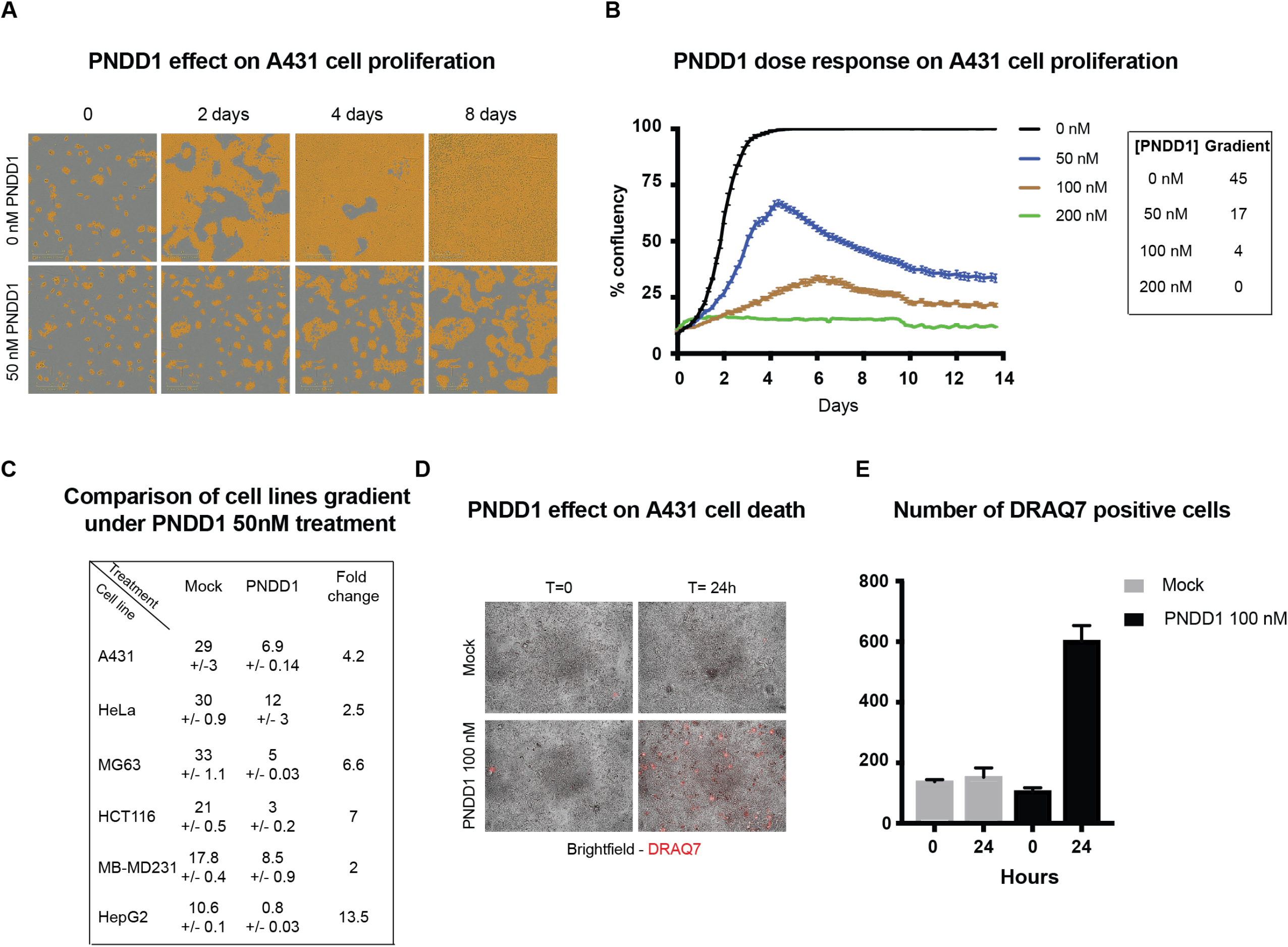
Different cancer cell lines are sensitive to PNDD. (**A**) Cell proliferation under PNDD1 treatment. A431 cells were treated with PNDD1 and bright field acquisition was made every 4 hours. Cell covered areas are revealed by a yellow mask. Representative of 3 independent experiments. (**B**) PNDD1 dose response effect on cell proliferation. Left panel: A431 cells were treated with 50, 100 and 200nM PNDD1 over 2 weeks. Bright field acquisition was made every 4 hours. A mask was used to identify areas covered by cells and measure confluency. Error bars at s.d. Representative of 3 independent experiments. Right panel: summary table of A431 gradient on the first 48 hours upon PNDD1 treatment at different concentrations. (**C**) Summary table of PNDD1 effect on different cell lines growth during 48 hours of treatment. (**D**) A431 were treated with or without 100 nM PNDD1 for 24 hours in presence of cell death marker DRAQ7. Brightfield and far red acquisition were made every 4 hours. Representative of 3 independent experiments. (**E**) Quantification of DRAQ7 positive A431 cells with (black bars) or without (grey bars) 100 nM PNDD1 treatment after 24 hours. Error bars at s.d. Representative of 3 independent experiments.

To evaluate if the effect of PNDD1 is widely applicable, we treated HeLa, HepG2 and MB-MDA231 cells with 50nM PNDD1, using the same live proliferation analysis method and using either PNDD0 or PBS as a control. All tested cell lines showed a decrease of cell proliferation, albeit with different sensitivities (Supplementary Fig. S4A-F). To compare the sensitivity of each cell line to the inhibitor, the gradient of confluency change over time was measured to estimate a growth rate. The different cell lines were exposed to 50nM PNDD1 (Fig. 4C). The HepG2 cell line showed the highest sensitivity with no cell proliferation at 50nM PNDD1 and the largest decrease in growth rate at 13.5-fold (Fig. 4C, Supplementary Fig. S4A). The effect in HeLa and MB-MDA-231 cells was approximately a 2-fold decrease in growth rate (Fig. 4C, Supplementary Fig. S4B, S4D).

To confirm the induction of cell death, we added DRAQ7, a far-red fluorescent dye that marks dead cells while incubating A431 with 100 nM PNDD1. A significant increase of DRAQ7 staining was apparent after 24h (Fig. 4D-E). This is consistent with the decrease of Renilla luciferase reporter signal observed after 24 hours of PNDD1 treatment (Fig. 3C, D). Overall, these results indicate that PNDD1 slows cell proliferation in many cancerous cell lines and can induce cell death of A431 cells if c-Myc inhibition is marked and sustained for several hours.

### PNDD1 inhibits Diffuse Large Cell Lymphoma proliferation

While c-myc is presumably involved in many different cancers, its implication in lymphomas has been well established. Diffuse Large B Cell Lymphomas (DLBCL), in particular, typically overexpress c-myc^17,18^. They are also difficult to treat. We obtained from clinicians several patient derived DLBCL lines: HT, OCI-LY3, OCI-LY19 and SUDHL2 and validated PNDD1 uptake and nuclear enrichment by cell fractionation (Fig. 5A, Supplementary Fig. S5A-B).

**Figure 5:**
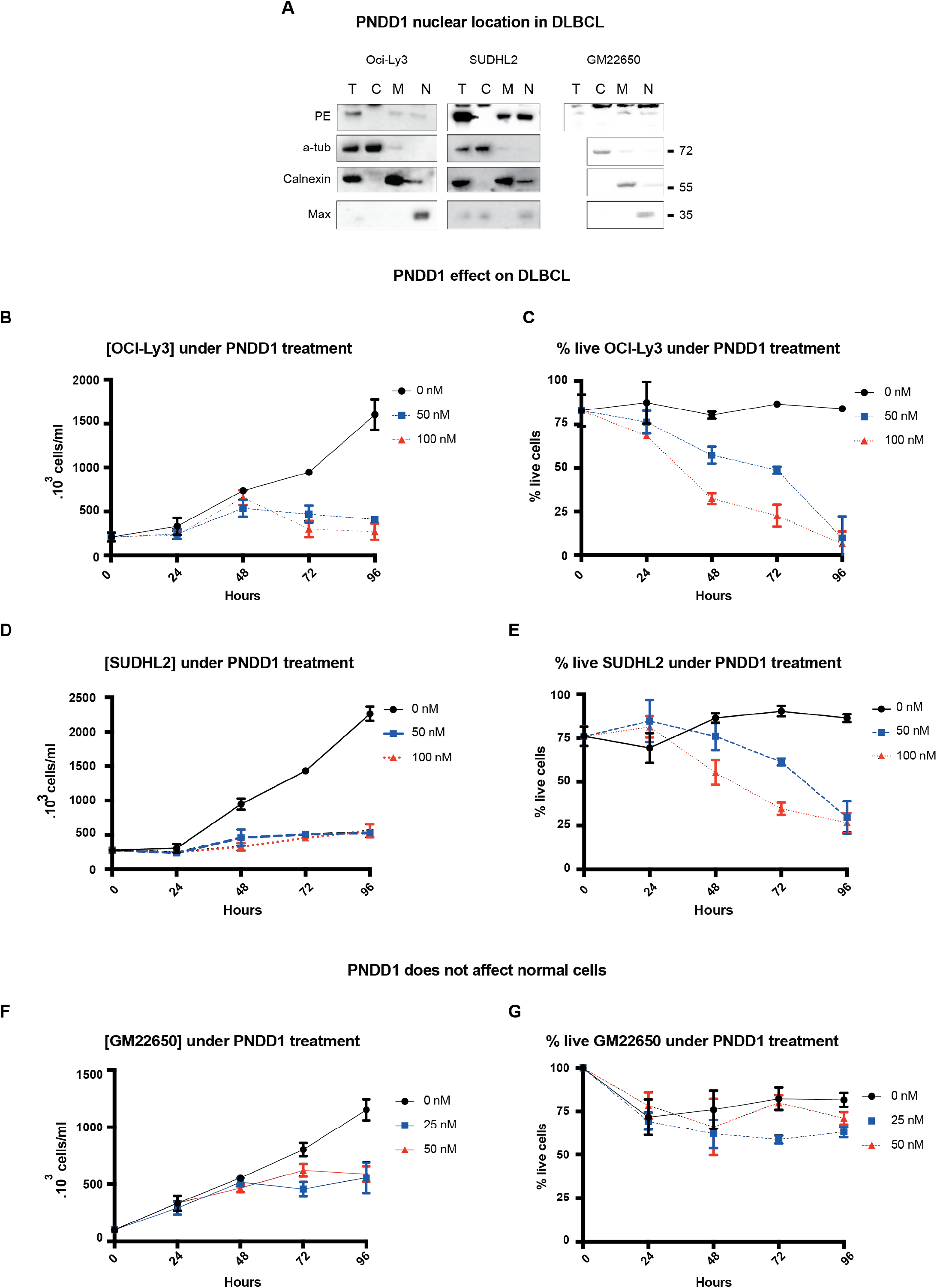
PNDD1 reduces DLBCL proliferation and induces cell death. (**A**) PNDD1 subcellular cell fractionation of OCI-Ly3, SUDHL2 and GM22650 cells. OCI-Ly3 (**B**-**C**) and SUDHL2 (**D**-**E**) cells were incubated with 50 or 100 nM PNDD1 for 4 days. (**B** and **D**) Cell density was measured every 24 hours. (**C** and **E**) Percentage of live cells were counted every 24 hours. Error bars at s.d. Representative of 3 independent experiments. (**F**) GM22650 cells were incubated with 25 or 50 nM PNDD1 for 4 days. Cell density was measured every 24 hours. (**G**) Percentage of live cells was counted every 24 hours. Error bars at s.d. Representative of 3 independent experiments.

Next, we incubated the DLBCL cell lines with 50 or 100 nM PNDD1 for 4 days. We observed a decrease in cell count across all the DLBCL cell lines 2 days after treatment in a dose dependent fashion (Fig. 5B, D, Supplementary Fig. S5C, E). The percentage of live cells also decreased in a dose dependent manner, indicating that the treatment induced cell death (Fig. 5C, E, Supplementary Fig. S5D, F). As expected, the effect was specific to H1 as PNDD0 had no similar effect (Supplementary Fig. S5G-H).

Among the cell lines, we could not observe a clear correlation between the amount of uptake of PNDD1 and reduction of cell proliferation. For instance, PNDD1 uptake is lower in OCI-LY3 than in SUDHL2 yet both cell lines displayed a similar rate of inhibition (Fig. 5C, E). We did, however, observe a clear difference with the immortalized B cell line GM22650, which displayed an arrest in proliferation but no cell death (Fig. 5F-G).

Overall, our results indicate that DLBCL would be a therapeutic indication for PNDD1 and that toxicity could be limited.

## Discussion

In this report, we show that a truncated toxoid form of PE can be used as a nuclear delivery device for biologics. Multiple protein toxins have evolved to efficiently penetrate cells to reach cytosolic or nuclear targets^7^. For this, they bind cell surface receptors that traffic to a compartment where the toxin can interact with a translocation machinery. PE binds to LRP1 and traffics to the ER before PE or its domain III is translocated to the cytosol and reaches its main target, the ribosome^7,8^. However, a fraction of LRP1 and thus PE also traffics to the NAE and the nucleus^2^. While this trafficking is quantitatively limited and seems to play only a minor role in PE intoxication, it is sufficient for PNDD to deliver cargo to the nucleus.

The PNDD approach is amenable to cargo of different sizes as demonstrated by the nuclear delivery of the 2 kDa H1 peptide, the 10 kDa Omomyc polypeptide and the 25 kDa GST protein. Thus, using various peptides, targeting a range of nuclear proteins implicated in cancer appears possible. However, the amount of chimeric protein delivered to the nucleus could represent a limitation. Indeed, our data of cell fractionation and immunofluorescence suggests that less than 10% of PNDD1 entering the cell reaches the nucleoplasm.

The transcription factor c-Myc is a key regulator of cell growth and its expression levels are tightly regulated^30^. This might explain why its inhibition by PNDD1 or 2 is working so efficiently. In fact, nanomolar doses of PNDD1/2 were enough to significantly reduce c-Myc activity and induce cell death in DBCL cell lines. This range is several orders of magnitude lower than the tested CPPs, which require micromolar doses to be effective. A recent study has reported small molecules able to inhibit Myc/Max interaction, which also require low micromolar range^31^. This significant difference could stem from a combination of at least two factors. First, based on previous work, it seems likely that PNDDs translocate through membranes using cellular machinery such as the Sec61 translocon, which could be directional transport a priori more efficient than physico-chemical diffusion through membranes^2,32^. Second, once in the nucleoplasm, PNDD1/2 are unlikely to diffuse out in the cytosol given their size, by contrast CPPs need to diffuse in the cytosolic fraction in addition to the nucleoplasm.

The reliance on the NAE pathway for delivery may also have a significant impact on the therapeutic index of PNDDs. Previous studies have reported that tumor cells tend to accumulate growth factor receptors in the nucleus, suggesting a more active NAE pathway^3,33^. It suggests that cancer cells may transfer more PNDDs to the nucleus than normal cells, impacting favorably the therapeutic index. Thus, the PNDD approach may be well suited for cancer treatment.

The therapeutic index of a treatment depends on many factors. It seems that tumor cells may be more sensitive than their normal counterpart to c-Myc inhibition, a phenomenon described as oncogene addiction^19^. The precise mechanism underlying cell death of cancer cells after c-Myc inhibition is not known. We observe a significant delay between decrease in Firefly luciferase activity and the onset of cell death: reduction of luciferase activity occurs within 4h of continuous incubation and is 80% reduced by 8h. In contrast, we observe cell death as well as a general inhibition of transcription after 24-48h in A431 and in DCBL cell lines. The Firefly luciferase assay relies on an mRNA with a 1.5 hours half-life and its protein product a 2-3 hour half life^34,35^. By contrast, it seems to take about 24 hours for Myc target genes to experience significant reduction at the protein level and induce a shutdown of cellular functions.

Obviously, the requirement for a lasting inhibition of c-Myc activity may be a challenge for the use of PNDD1/2. Given the 4h intracellular half-life of PNDD1, a constant delivery over 24 hours may be required to irreversibly commit cancer cells to apoptosis. Alternatively, our data shows that it is possible to increase the intracellular half-life of PNDDs by treating cells concomitantly with a proteasome inhibitor such as bortezomib. Since bortezomib is also used sometimes for lymphoma treatment, it seems possible to combine it with PNDD type molecules for therapeutic purposes.

A potential therapeutic indication could be the lymphomas of the DLBCL type. PNDD1 was efficient to slow down cell proliferation and induce cell death in all the patient cell lines we tested. Even in cell lines with comparatively lower PNDD1 uptake, such as OCI-Ly3, the induction of cell death was robust. It might be because these cells are highly dependent on Myc activity, as a study has recently shown that OCI-LY3 contains multiple copies of MYC^36^. By contrast, the immortalised normal B-cells were able to stop proliferation without significant cell death.

In conclusion, PNDD appears to be an efficacious method to deliver peptidic cargo in the nucleus, with potentially many different applications. In the examples we tested, PNDD did not affect the functional properties of its cargos, H1 and Omomyc. It is efficacious at much lower concentrations than Cell Penetrating Peptides that have demonstrated therapeutic abilities^37^. Further characterisation and optimisation of PNDD1/2 might be required to obtain serum and intracellular half-lives values that would be compatible with in vivo use.

## Material and Methods

### Gene synthesis and cloning

The sequences of PNDD0 and PNDD1 were synthesized and cloned in a pET100/D-TOPO vector using gene synthesis (Thermo Fisher Scientific, Carlsbad, CA). H1 peptide sequence is underlined.

PE151: MEEAFDLWNECAKACVLDLKDGVRSSRMSVDPAIADTNGQGVLHYSMVLEGGNDALKLAIDNALSITSDGLTIRLEGGVEPNKPVRYSYTRQARGSWSLNWLVPIGHEKPSNIKVFIHELNAGNQ

PE212: MEEAFDLWNECAKACVLDLKDGVRSSRMSVDPAIADTNGQGVLHYSMVLEGGNDALKLAIDNALSITSDGLTIRLEGGVEPNKPVRYSYTRQARGSWSLNWLVPIGHEKPSNIKVFIHELNAGNQLSHMSPIYTIEMGDELLAKLARDATFFVRAHESNEMQPTLAISHAGVSVVMAQAQPRREKRW

PE277: MEEAFDLWNECAKACVLDLKDGVRSSRMSVDPAIADTNGQGVLHYSMVLEGGNDALKLAIDNALSITSDGLTIRLEGGVEPNKPVRYSYTRQARGSWSLNWLVPIGHEKPSNIKVFIHELNAGNQLSHMSPIYTIEMGDELLAKLARDATFFVRAHESNEMQPTLAISHAGVSVVMAQAQPRREKRWSEWASGKVLCLLDPLDGVYNYLAQQRCNLDDTWEGKIYRVLAGNPAKHDLDIKPTVISHRLHFPE

PE389/PNDD0: MEEAFDLWNECAKACVLDLKDGVRSSRMSVDPAIADTNGQGVLHYSMVLEGGNDALKLAIDNALSITSDGLTIRLEGGVEPNKPVRYSYTRQARGSWSLNWLVPIGHEKPSNIKVFIHELNAGNQLSHMSPIYTIEMGDELLAKLARDATFFVRAHESNEMQPTLAISHAGVSVVMAQAQPRREKRWSEWASGKVLCLLDPLDGVYNYLAQQRCNLDDTWEGKIYRVLAGNPAKHDLDIKPTVISHRLHFPEGGSLAALTAHQACHLPLETFTRHRQPRGWEQLEQCGYPVQRLVALYLAARLSWNQVDQVIRNALASPGSGGDLGEAIREQPEQARLALTLAAAESERFVRQGTGNDEAGAAS

PNDD1: MEEAFDLWNECAKACVLDLKDGVRSSRMSVDPAIADTNGQGVLHYSMVLEGGNDALKLAIDNALSITSDGLTIRLEGGVEPNKPVRYSYTRQARGSWSLNWLVPIGHEKPSNIKVFIHELNAGNQLSHMSPIYTIEMGDELLAKLARDATFFVRAHESNEMQPTLAISHAGVSVVMAQAQPRREKRWSEWASGKVLCLLDPLDGVYNYLAQQRCNLDDTWEGKIYRVLAGNPAKHDLDIKPTVISHRLHFPEGGSLAALTAHQACHLPLETFTRHRQPRGWEQLEQCGYPVQRLVALYLAARLSWNQVDQVIRNALASPGSGGDLGEAIREQPEQARLALTLAAAESERFVRQGTGNDEAGAAS<u>NELKRAFAALRDQI</u>

### Cell line culture and stable transduction

All cell lines come from ATCC. WT A431 cells were transduced with Cignal Lenti Myc Reporter (Qiagen, USA) expressing Firefly luciferase under E-Box promoter and CMV-Renilla luciferase under CMV promoter. Single clone was selected after colony isolation and Firefly/Renilla luciferases expression level tested. A431, HepG2, MG63, HeLa, HCT116 and MB-MDA231 were maintained in high-glucose Dulbecco’s modified Eagle’s medium supplemented with 10% foetal calf serum (FBS). OCI-Ly3, OCI-Ly19, SUDHL2 and HT cells were maintained in Roswell Park Memorial Institute (RPMI) 1640 supplemented with 20% FBS. GM22650 cells were maintained in RPMI 1640 supplemented with 15% FBS. All cell lines were maintained at 37°C in a 10% CO2 incubator. All experiments were performed on cells passaged fewer than 10 times after thawing.

### Firefly luciferase and Renilla luciferase activity reading

40,000 A431-mrep cells were seeded in a 96-well plate (Falcon) 48 hours. Luminescence is detected using the Promega Dual-Glo luciferase assay system, according to the manufacturer’s protocol and read using Tecan Infinite M200 microplate reader using 100 ms integration time.

### Bacterial expression

Purified plasmids previously described were introduced into the E. coli strain Epicurean BL21 (Stratagene, USA). Cultures were grown at 37°C until OD A600 value reached 0.5 before the induction of protein expression by addition of Isopropyl *β* -D-1-thiogalactopyranoside (0.1 mM) in the LB culture medium. After 2 hours of induction at 37°C, bacteria were recovered by centrifugation for 30 minutes at 3000 g at 4°C.

### Purification of PNDD0, PNDD1 and PNDD2

Bacteria expressing PNDD0, PNDD1 or PNDD2 were resuspended in 3 mL of lysis buffer (Tris 50 mM pH 8, NaCl 170 mM, imidazole 20 mM, urea 6 M, NP40 0.5% v/v) for a pellet corresponding to 60 mL bacterial culture. The solution was homogenized and sonicated. Insoluble material was discarded by centrifugation for 30 minutes at 16,000 g at 4°C. 2ml of packed Ni-NTA affinity chromatography resin (Qiagen, USA) was calibrated with 10 volumes of lysis buffer, before being incubated with soluble PNDD0, PNDD1 or PNDD2 overnight on rotor at 4°C. Resin was then washed with 10 volumes of washing buffer (Tris 50 mMpH 8, NaCl 170 mM, imidazole 40 mM, urea 6 M, NP40 0.5% v/v). PNDD0, PNDD1 or PNDD2 were eluted twice with 1 volume of elution buffer (Tris 50 mM pH 8, NaCl 170 mM, imidazole 1 M, urea 6 M, NP40 0.5% v/v). Purified protein elute was injected in Slide-A-lyzer Dialysis Cassette 20,000 MWCO (Thermo Scientific) and incubated in 3 baths of 100 volumes of PBS for 21 hours at 4°C. Dialysed elute is retrieved from the cassette and tested for solubility by spinning down for 30 minutes at 15,000g 4°C. The supernatant is retrieved and stored at −80°C. The pellet is resuspended in the same volume with lysis, washing or elution buffer. A sample is taken from each fraction to compare the amount of purified protein.

### PNDD0 and PNDD1 quantification and quality control

Purified protein quantification was done by Bradford by comparing the optical density at 595 nm (OD595) nm with a BSA standard. Purity control was made by running samples on SDS-PAGE followed by coomassie blue staining (InstantBlue™ Protein Stain, Expedeon). For experiments in Figure 4 and 5; we verified the quality and activity of new batches of PNDD0 and 1 using the following procedures. Testing uptake in MG63 cells with incubation at 500 nM for 1 hour followed by immunofluorescence with anti-PE antibody to test presence in NAE. Anti-Myc Activity testing by incubation at concentrations of 10 to 100 nM using an A431-mrep cells. For validation, PNDD1 EC50 must be around 25 nM range at 6 hours incubation at 37°C and no significant effect on renilla. PNDD0 must have a non-significant effect on either luciferases.

### Immunofluorescence

PE (#341215, Merck Millipore, Darmstadt, Germany) was labelled with AnaTag Hilyte™ Fluor 594 and purified truncated PE constructs were labelled with AnaTag Hilyte™ Fluor 488 according to the manufacturer’s instructions (#72048, AnaSpec, Fremont, CA). Cells were seeded onto glass coverslips in 6-well dishes (Thermo Fisher Scientific) and incubated at 37 °C, 10% CO2 for 8–16 h before PE treatment at a concentration of 0.5uM for 1h. All subsequent steps were performed at room temperature. Cells were washed with Dulbecco’s Phosphate Buffer Saline (D-PBS), fixed for 15 min using 4% paraformaldehyde 2% sucrose in D-PBS, washed with D-PBS once, then permeabilized and blocked with 0.2% Triton X-100, 2% Foetal Bovine Serum (FBS) in D-PBS for 15 min. Cells were washed three times using D-PBS and then subsequently stained for 45 min Hoechst 33342 (Invitrogen) at a concentration of 1ug/ml. Cells were mounted onto glass slides using FluorSave (Merck Millipore).

### Cell fractionation

Cells were seeded at the desired densities in six-well dishes (Thermo Fisher Scientific) and incubated at 37°C, 10% CO2 for 16 hours. Cells were treated for 1 hour with mock (untreated cells) or PNDD1 200 nM before fractionation with Subcellular Protein Fractionation Kit (#78840, Thermo Fisher Scientific) according to the manufacturer’s protocol. In addition, cytosolic fraction is centrifuge at 15,000g for 15 minutes to remove membrane contaminants. Samples are denaturated in Laemmli blue 2X and heat-denaturated for 5 minutes at 95°C. Samples are loaded on SDS-PAGE followed by western blot.

### Cell proliferation after PNDD1 treatment

Cells were seeded at 10 to 20% confluency. 24 hours later, the cells were incubated with different doses of PNDD1 at 37°C, 10% CO2 for 14 days. Live imaging was performed using IncuCyte ZOOM^®^ Live-Cell Analysis System. Bright field Images were taken every 6 hours. Images were analysed using IncuCyte ZOOM^®^ Live-Cell Analysis software.

### DRAQ7 assay

25000 HepG2 cells were seeded per well in a 96-well plate. After incubation at 37°C for 24 hours, DRAQ7 dye was prepared with 100 nM PNDD1 at 500 times dilution. Cell mortality was tracked by acquiring nine fields of brightfield and far-red images at 20x magnification every 4 hours using the Operetta machine (Perkin-Elmer) for 3 to 5 days. The machine kept the plate incubated at 37°C with 8% CO2.

### Quantitative RT-PCR (qRT-PCR) after PNDD1 treatment

400000 A431 cells were seeded in each well in a 6 well plate (Falcon) 24 hours before incubation with 100 nM PNDD1 for 16 hours at 37°C. Total RNA was isolated using the RNeasy Mini Kit (Qiagen, Ref. no 74106) as described by manufacturer’s protocol. 20μg total RNA was reversed transcribed using SuperScript III First-Strand Synthesis System for RT-PCR (Invitrogen, Cat. no. 18080051) as described by manufacturer’s protocol. The relative expression of mRNAs of c-Myc regulated genes identified in RT^2^ Profiler™ PCR Array Human MYC Targets (Qiagen, Cat. no. PAHS-177Z) was determined by real-time quantitative RT-PCR. Briefly, a PCR component mix containing cDNA, RT2 SYBR Green Mastermix (Qiagen, Cat. no. 330523) and RNase-free water is prepared. 25 μl of the PCR reaction was added to each well in one array plate and subjected to a qPCR program using the ABI 7500. PCR cycling conditions consisted of an initial denaturation step at 95°C for 10 minutes followed by an amplification program for 40 cycles of 15 seconds at 95°C, and 60 seconds at 60°C with fluorescence acquisition at the end of each extension. The relative expression of each gene between treated and untreated samples is calculated using the comparative ΔΔCT method, using the mock treated sample as calibrator and housekeeping gene as internal control.

### DLBCL cell proliferation and cell death assay

200,000 to 300,000 cells were seeded 24 hours before PNDD0 or PNDD1 50 nM treatment. Three samples of the suspension cell culture was taken every 24 hours after PNDD0 or PNDD1 50 nM treatment to measure the cell number and the percentage of dead cells. Three aliquots of supernatant were taken and mixed with equal volume of Trypan blue before reading on Countless Cell Automated Cell Counter^®^ (Invitrogen).

## Acknowledgement

We thank Pr Anand Jeyasekharan (NUH, Singapore) and Dr Widanalage Sanjay Prasad De Mel (NUH, Singapore) for scientific discussion and providing DLBCL cell lines. This work was funded by A.C. BMRC Young Investigator Grant and the Agency for Science, Technology and Research.

## Author contributions

A.C and F.B. designed the study. A.C and T.A.T. performed experiments. T.A.T., A.C and F.B. wrote the manuscript.

## Disclosure

PNDD1/tPE-H1 method to inhibit c-Myc is filled under the title and patent application:

### Chimeric Molecule for Targeting c-Myc in Cells

Patent Application No PCT/SG2018/050584.

## Supplementary figure legends

**Figure S1: PNDD purification.**

**A-F** PE389 purification; **G-H** PE277 purification; **I-J** PE212 purification; **K-L** PE151 purification. In: Input, total bacterial extract; Out: Output, unbound proteins; Sn: Supernatant; P: Pellet; Et: Elution total, E1+E2.

**Figure S2: PNDD1 characterisation.**

PNDD1 solubility: (**A**) PNDD1 purification followed by SDS-PAGE stained with coomassie blue. T: Total, Sn: supernatant, P: Pellet, Out: output, Et: Elution total. Molecular weights are shown on the right. (**B**) Coomassie staining replicates showing PNDD1 solubility after dialysis and spin down. E: Elute before dialysis and centrifugation; Sn: Supernatant after centrifugation; P: Pellet after centrifugation. Molecular weights are shown on the right (left panel). PNDD1 subcellular distribution: (**C**) MG63 cell fractionation after 1 hour treatment with PNDD1. Alpha-tubulin (A-Tub), calnexin and MAX are used as fraction controls for respectively C: Cytosolic fraction; M: Membrane fraction; N: Nuclear fraction. Representative western blot of 3 independent experiments. (**D**) Left panel: PNDD1 cell fractionation. Quantification of ratio nucleus/total extract (under corresponding band). PNDD1 N/T average ratio = 0.14. C: Cytosolic fraction; M: Membrane fraction; N: Nuclear fraction. Antibodies are labelled on the left of each blot. Molecular weight is shown on the right. Middle panel: Max cell fractionation. Quantification of ratio nucleus/total extract (under corresponding band). Max N/T average ratio = 2.2 C: Cytosolic fraction; M: Membrane fraction; N: Nuclear fraction. Antibodies are labelled on the left of each blot. Molecular weight is shown on the right. Normalisation PE nuclear proportion over Max: 0.14/2.2= 6.3%. Right panel: table summarizing PNDD1 N/T ration, Max N/T ration and PNDD1 normalisation over Max. PNDD1 stability: (**E**) PNDD1 uptake kinetic during continuous incubation. Total cell lysates were lysed at different time points up to 30 hours and PNDD1 remaining amount was analysed by western blot to determine its half life in cells. EF2 was used as a loading control. Molecular weights are shown on the right. (**F**) Replicates 1 and 2 of Fig. 2B. PNDD1 was incubated 1 hour (pre-) with MG63 cells before washing with PBS and replacing with media (post-). Total cell lysates were lysed at different time points up to 30 hours and PNDD1 remaining amount was analysed by western blot to determine its half life in cells. EF2 was used as a loading control. Molecular weights are shown on the right. Representative western blot of 3 independent experiments. (**G**) Replicates of Fig. 2E. MG63 cells were incubated with PNDD1 during 1 hour with or without Bortezomib before wash of PE. As a control, MG63 cells were incubated continuously during 24 hours with PNDD1. EF2 was used as a loading control. Molecular weights are shown on the right. Triplicate experiments are shown.

**Figure S3: PNDD1 and PNDD2 effect on c-myc controlled gene expression**

(**A**) A431 cell fractionation after 1 hour PNDD1 treatment. A-Tub, calnexin and Max are used as fraction controls for respectively C: Cytosolic fraction; M: Membrane fraction; N: Nuclear fraction. Antibodies are labelled on the left of each blot. Molecular weight is shown on the right. Representative western blot of 3 independent experiments. (**B**) PNDD1 dose response effect on E-Box Firefly luciferase and CMV Renilla luciferase after 6 hours incubation. (**C**) Replicates of A431 cells expressing Firefly luciferase (black bars) and Renilla (grey bars) under CMV promoter control treated for 6 hours with 50 nM PNDD0. Error bars are s.d. (**D**) A431 cell fractionation after 1 hour PNDD2 treatment. A-Tub, calnexin and Max are used as fraction controls for respectively C: Cytosolic fraction; M: Membrane fraction; N: Nuclear fraction. Antibodies are labelled on the left of each blot. Molecular weight is shown on the right. Representative western blot of 3 independent experiments. (**E**) PNDD2 dose response effect on E-Box Firefly luciferase activity after 6 hours continuous incubation. Representative of 3 independent experiments. (**F**) Firefly luciferase activity at different time points after continuous incubation (solid line) or after 1 hour treatment followed by a wash (dotted line). Error bars at s.d. (**G**) Replicates of A431 cells treated with PNDD1 overnight before RNA extraction. mRNA transcript levels of genes regulated by c-Myc were quantified by RT-PCRQ and compared with or without PNDD1 treatment. Housekeeping mRNAs (HPRT1, GAPDH) whose expression are not regulated by c-Myc are analysed in the same manner. Y axis shows the mRNA log2(fold change). Upregulated genes appear with negative log2 (fold change) compared to housekeeping genes. Results are significant below −1.5. Error bars at s.d. (**H**) Replicate of CPP-H1 effect on Firefly luciferase activity. Error bars at s.d. (**I**) Comparative optimum to obtain 80% Firefly luciferase activity decrease after 6 hours treatment with cell targeting peptides (CPP) fused to H1 and PNDD1 on A431 cells expressing Firefly luciferase under E-Box promoter control. X axis is shown in Log Cadherin (CAD; LLIILLRRRIRKQAHAHSK;) EC_50_=75 mM, Antenapedia (Int; RQIKIWFQNRRMKWKK) EC_50_=200 mM and TAT (GRKKRRQRRRPPQ) EC_50_=500 mM. Error bars at s.d.

**Figure S4**

Cell lines sensitivity to PNDD1. Comparison of PNDD1 50 nM effect on cell proliferation of (**A**) HepG2, (**B**) HeLa, (**C**) A431, (**D**) MB-MDA231 cells. Effect of PNDD1 50 nM on cell proliferation of (**E**) HCT116 and (**F**) MG63.

**Figure S5**

(**A**) PNDD1 cellular uptake in HT, Oci-Ly3, Oci-Ly19, SUDHL2 and MG63 (control) cells. EF2 was used as a loading control. Molecular weights are shown on the right. (**B**) PNDD1 subcellular cell fractionation of HT and Oci-Ly19 cells. Oci-Ly19 cells were incubated with 50 or 100 nM PNDD1 during 4 days. (**C, E, G**) Cell density was recorded every 24 hours. (**D, F, H**) percentage of live cells was recorded every 24 hours. Error bars are s.d. Representative of 3 independent experiments.

