## Supplementary figures for "Targeting c-Myc with a novel Peptide Nuclear Delivery Device"

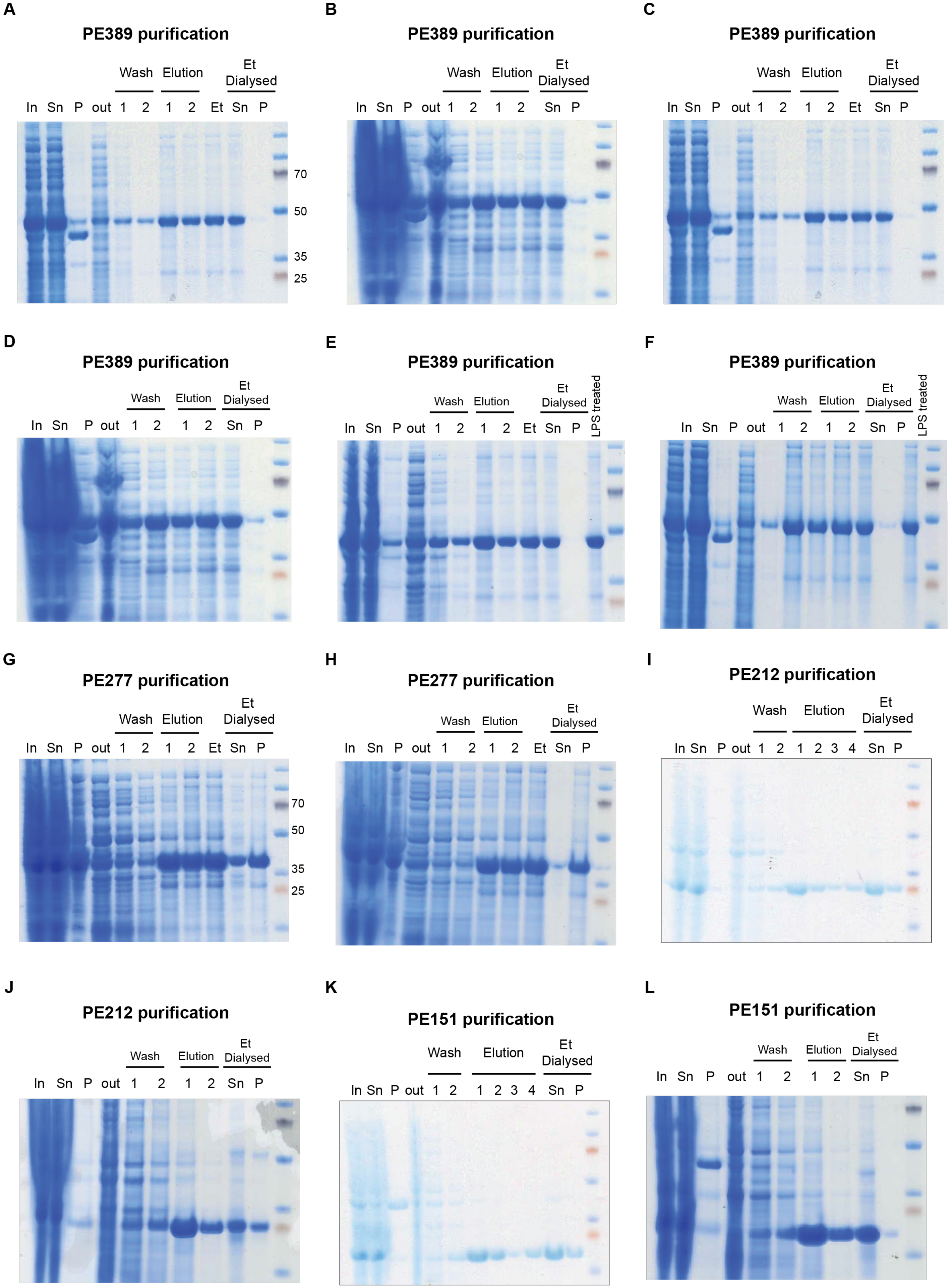


**Supplementary figure 1**

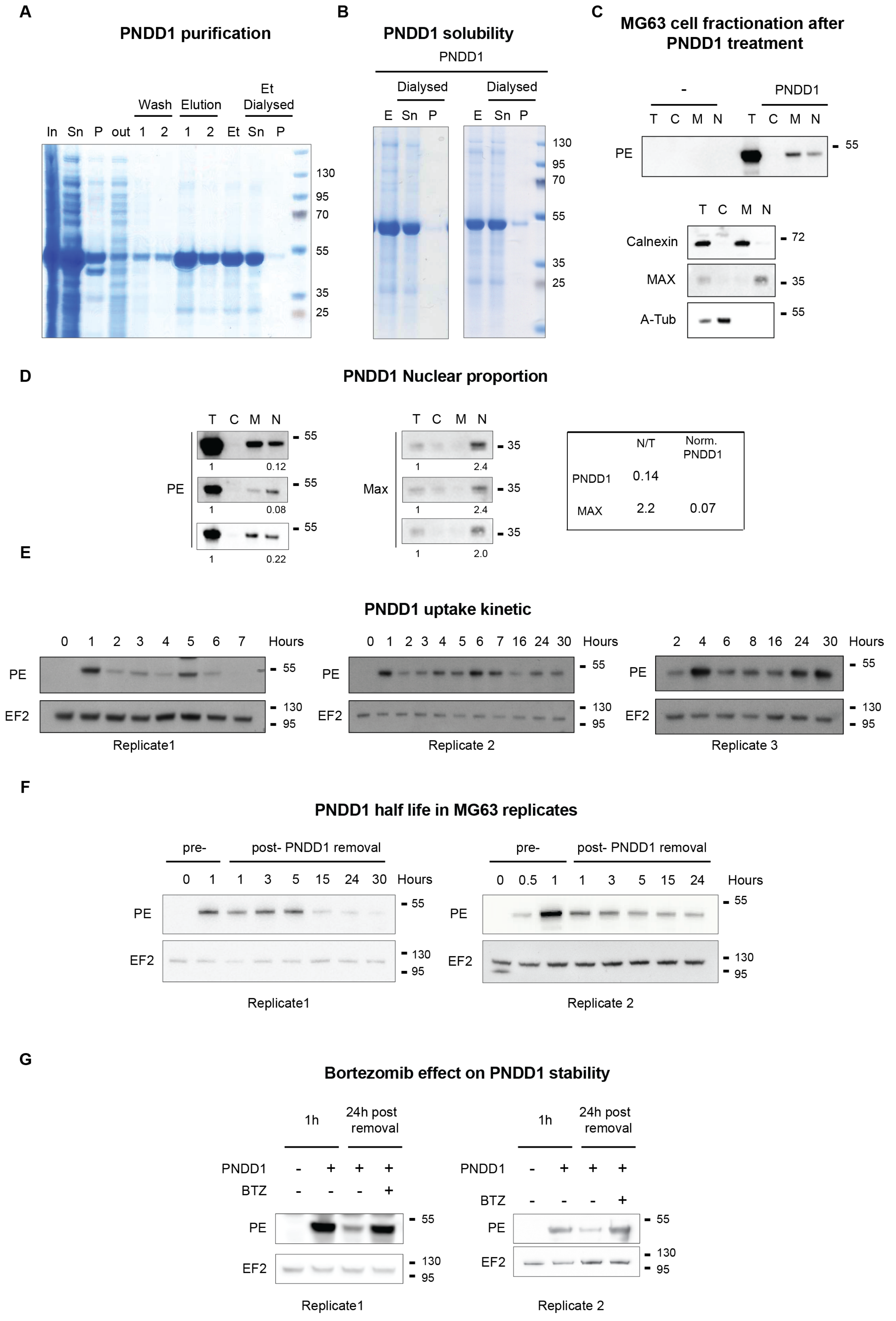


supplementary figure 2

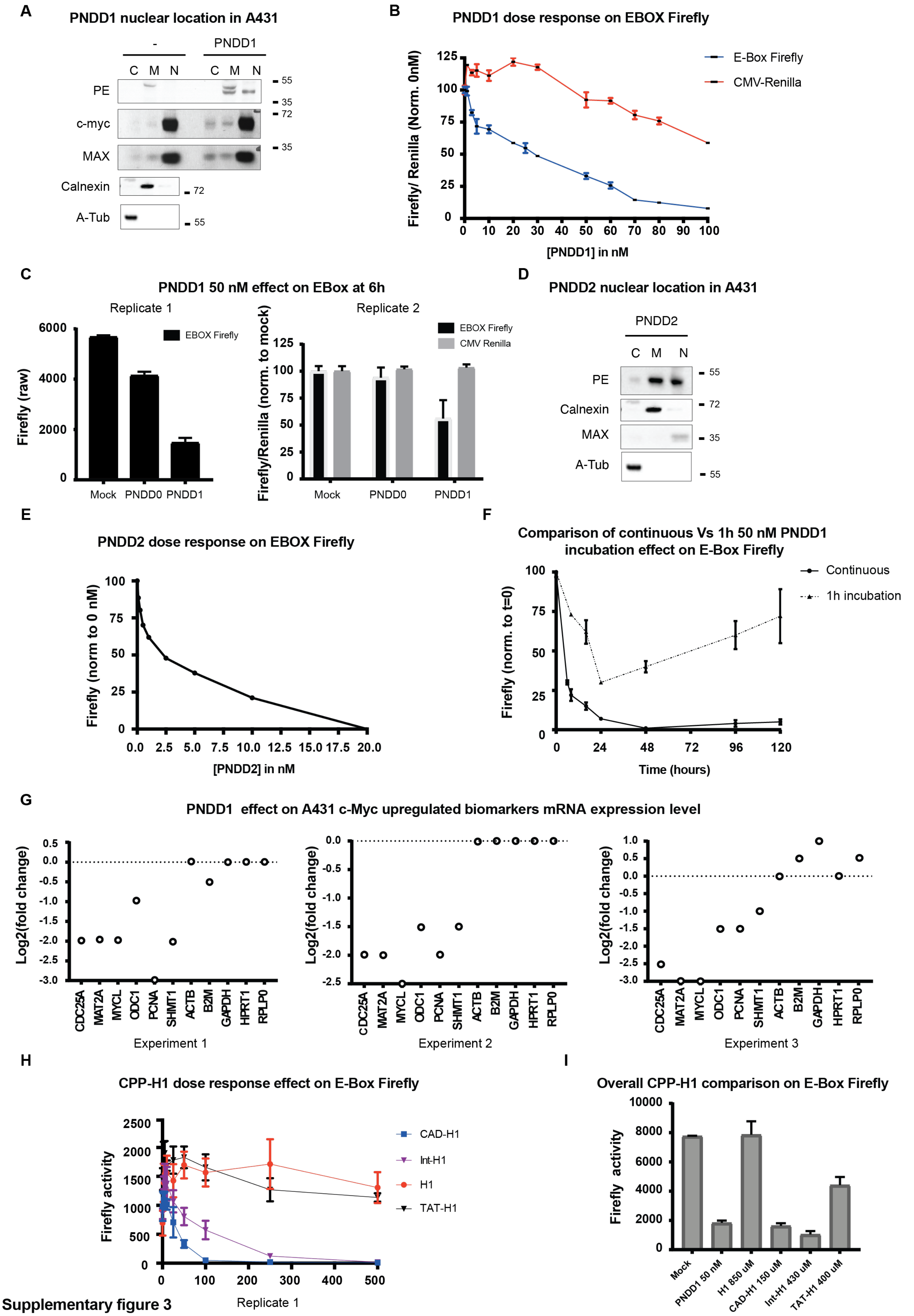


Supplementary figure 3

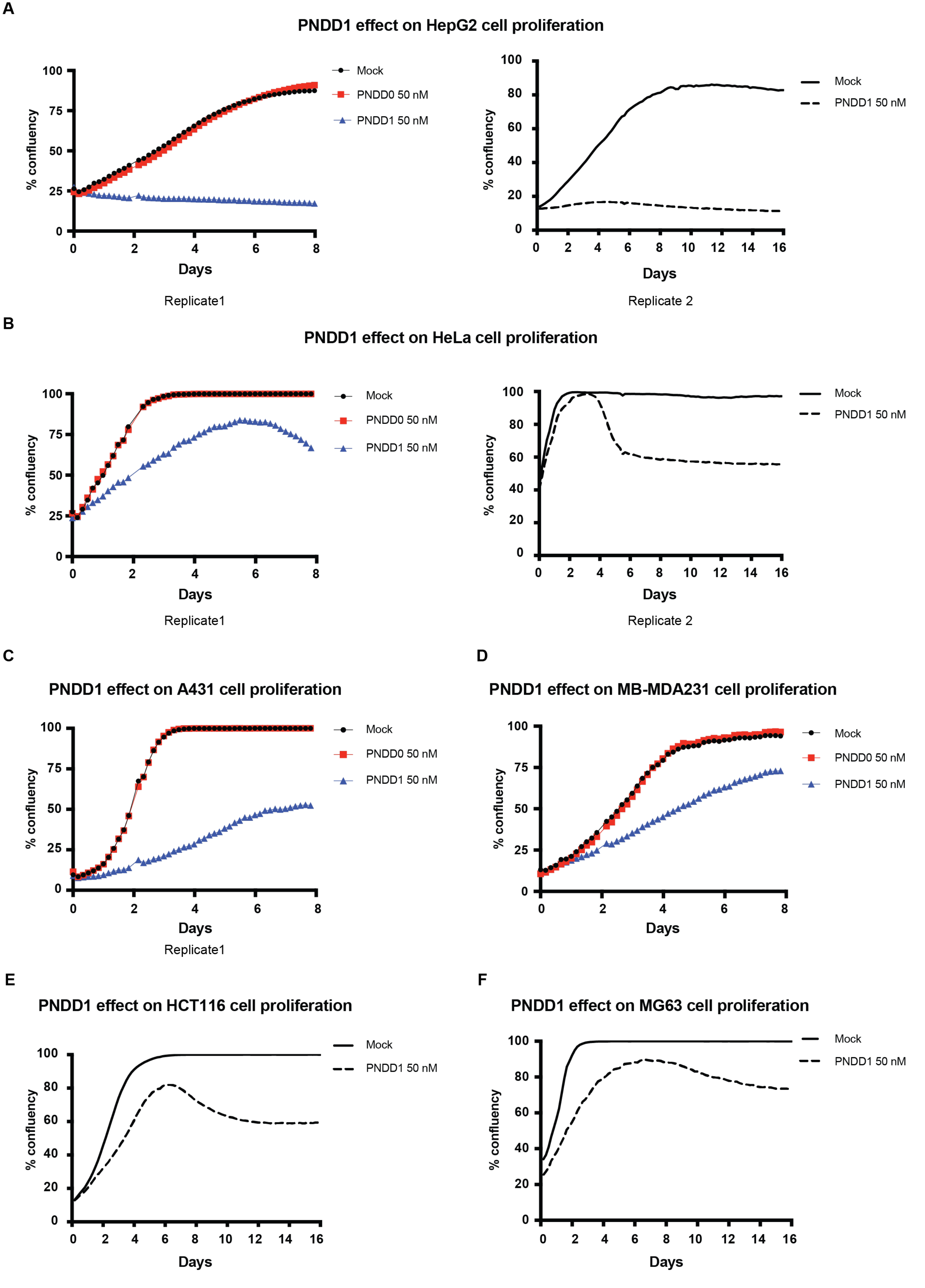


supplementary figure 4

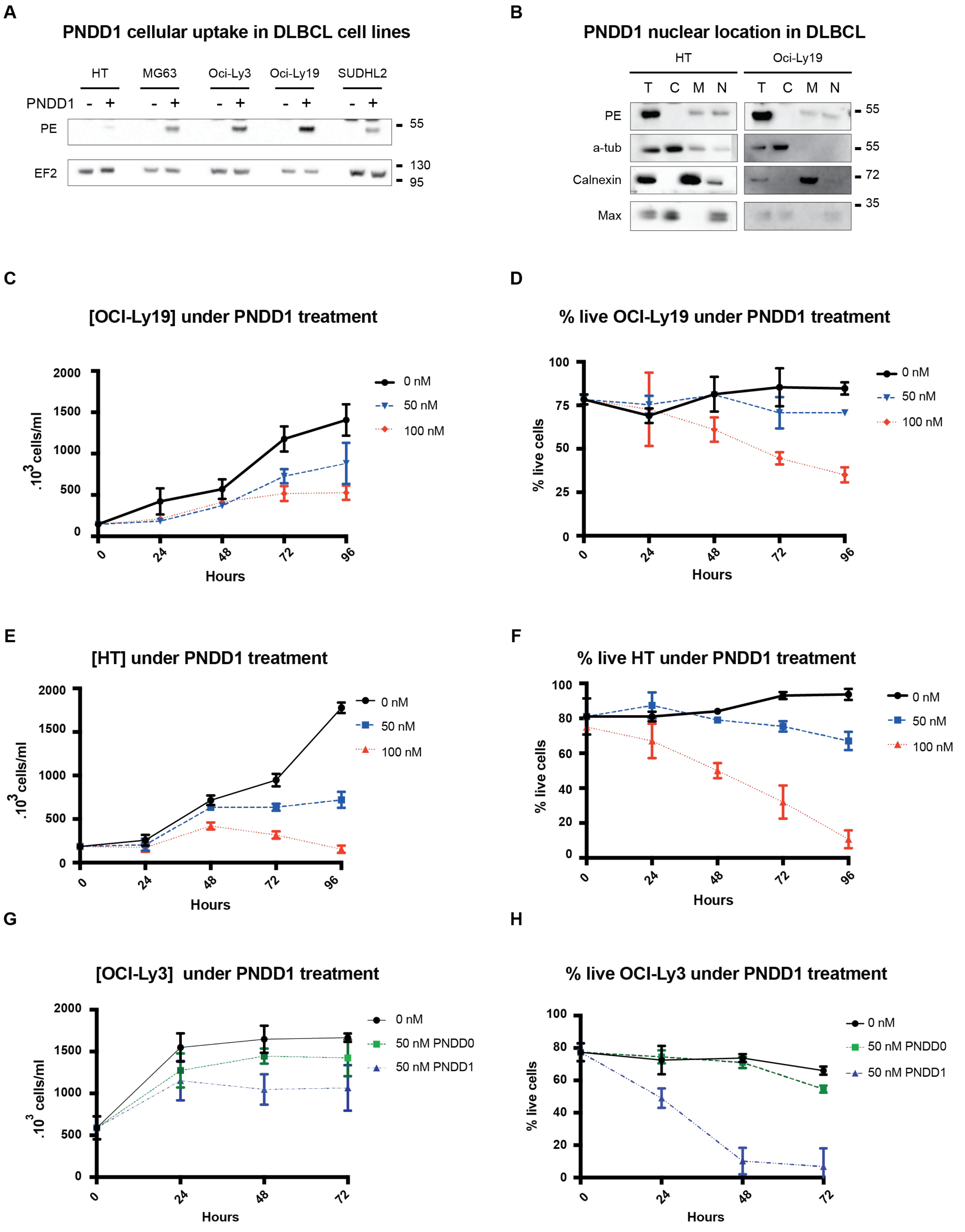


supplementary figure 5
